# p62-mediated Selective Autophagy Endows Virus-transformed Cells with Insusceptibility to DNA Damage under Oxidative Stress

**DOI:** 10.1101/502823

**Authors:** Ling Wang, Mary E. A. Howell, Ayrianna Sparks-Wallace, Caroline Hawkins, Camri Nicksic, Carissa Kohne, Kenton H. Hall, Jonathan P. Moorman, Zhi Q. Yao, Shunbin Ning

## Abstract

DNA damage response (DDR) and selective autophagy both can be activated by reactive oxygen/nitrogen species (ROS/RNS), and both are of paramount importance in cancer development. The selective autophagy receptor and ubiquitin (Ub) sensor p62 plays a key role in their crosstalk. ROS production has been well documented in latent infection of oncogenic viruses including Epstein-Barr Virus (EBV). However, p62-mediated selective autophagy and its interplay with DDR have not been investigated in these settings. In this study, we provide evidence that considerable levels of p62-mediated selective autophagy are constitutively induced, and correlates with ROS-Keap1-NRF2 pathway activity, in virus-transformed cells. Inhibition of autophagy results in p62 accumulation in the nucleus, and promotes ROS-induced DNA damage and cell death, as well as downregulates the DNA repair proteins CHK1 and RAD51. In contrast, MG132-mediated proteasome inhibition, which induces rigorous autophagy, promotes p62 degradation but accumulation of the DNA repair proteins CHK1 and RAD51. However, pretreatment with an autophagy inhibitor offsets the effects of MG132 on CHK1 and RAD51 levels. These findings imply that p62 accumulation in the nucleus in response to autophagy inhibition promotes proteasome-mediated CHK1 and RAD51 protein instability. This claim is further supported by the findings that transient expression of a p62 mutant, which is constitutively localized in the nucleus, in B cell lines with low endogenous p62 levels recaptures the effects of autophagy inhibition on CHK1 and RAD51 protein stability. These results indicate that proteasomal degradation of RAD51 and CHK1 is dependent on p62 accumulation in the nucleus. However, small hairpin RNA (shRNA)-mediated p62 depletion in EBV-transformed lymphoblastic cell lines (LCLs) had no apparent effects on the protein levels of CHK1 and RAD51, likely due to the constitutive localization of p62 in the cytoplasm and incomplete knockdown is insufficient to manifest the effects on its nuclear function. Furthermore, shRNA-mediated p62 depletion in EBV-transformed LCLs results in significant increases of endogenous RNF168-γH2AX damage foci and chromatin ubiquitination, indicative of activation of RNF168-mediated DNA repair mechanisms. Our results have unveiled a pivotal role for p62-mediated selective autophagy that governs DDR in the setting of oncogenic virus latent infection, and provide a novel insight into virus-mediated oncogenesis.

**Author Summary:** Reactive oxygen/nitrogen species (ROS/RNS) can induce both DNA damage response (DDR) and selective autophagy, which play crucial roles in cancer development. The selective autophagy receptor and ubiquitin (Ub) sensor p62 links their crosstalk. However, p62-mediated selective autophagy and its interplay with DDR have not been investigated in latent infection of oncogenic viruses including Epstein-Barr Virus (EBV). In this study, we provide evidence that p62-mediated selective autophagy is constitutively induced in virus-transformed cells, and that its inhibition exacerbates ROS-induced DNA damage, and promotes proteasomal degradation of CHK1 and RAD51 in a manner depending on p62 accumulation in the nucleus. However, rigorous autophagy induction results in accumulation of DNA repair proteins CHK1 and RAD51, and p62 degradation. Further, transient expression of a constitutive nucleus-localizing mutant of p62 recaptured the effects of autophagy inhibition on CHK1 and RAD51 protein stability. These findings support the claim that p62 accumulation in the nucleus in response to autophagy inhibition promotes proteasome-mediated CHK1 and RAD51 protein instability. However, small hairpin RNA (shRNA)-mediated p62 depletion did not affect CHK1 and RAD51 protein levels; rather, shRNA-mediated p62 depletion activates RNF168-dependent DNA repair mechanisms. Our results have unveiled a pivotal role for p62-mediated selective autophagy in regulation of DDR by overriding traditional DDR mechanisms in the setting of oncogenic virus latent infection, and provide a novel insight into the etiology of viral cancers.

## Introduction

p62 (also named EBIAP, ZIP3, SQSTM1/Sequestosome-1), a human homolog of mouse ZIPs (Zeta PKC-interacting proteins), is well known as a selective autophagy receptor and a ubiquitn sensor, which controls myraid cellular processes, including redox homeostasis, DNA damage response (DDR), cancer development, aging, inflammation and immunity, osteoclastogenesis, and obesity, with or without the involvement of autophagy [1–3].

Autophagy, with either non-selective (random) or selective fashion, is a unique intracellular process that engulfs damaged and even functional cellular constituents and delivers them to lysosomes for digestion and recycling in the cytosol under diverse stresses, such as nutrient deprivation, viral replication, cancer hypoxia, genotoxic stress, and replicative crisis. Autophagy is thereby a crucial cellular machinery conserved from yeast to higher eukaryotes that maintains organ metabolism, genome stability, and cell survival, and functions as either tumor suppressor at early stage or promotor at late stage [4–6]. Distinct from non-selective autophagy, selective autophagy sort specific substrates to lysosomes, and is mediated by an increasing pool of receptors, including p62, NBR1, TAX1BP1, NDP52, OPTN, TRIMs, and TOLLIP [3, 7–10].

Reactive oxygen/nitrogen species (ROS and RNS), the major cause of endogenous DNA damage, can be produced in chronic viral infections, in which viral replication is generally absent [11]. They can directly modify DNA and generate different levels of lesions, including double strand breaks (DSBs) [12, 13]. Eukaryotic organisms have developed sophisticated strategies to repair DNA damage to ensure genomic integrity, with homologous recombination (HR) and nonhomologous end joining (NHEJ) being two non-redundant repair mechanisms for DSBs [14]. Unrepaired DSBs, however, incite chronic inflammation, resulting in genomic instability that promotes malignant transformation under certain conditions [15].

ROS/RNS also induce p62 expression through the Keap1-NRF2 pathway, licensing the induction of p62-mediated selective autophagy [16]. Mounting evidence indicates that DDR and selective autophagy closely crosstalk in response to oxidative stress, in which p62 plays a key role [17]. While p62 inhibits DNA damage repair, p62-mediated selective autophagy promotes DNA repair by targeting ubiquitinated substrates including p62 itself for degradation in cancer cells [18, 19], which usually harbor deficient traditional DNA repair mechanisms and heavily rely on autophagy as an alternative repair mechanism for survival [20, 21]. In this sense, p62-mediated selective autophagy, which is activated upon DNA damage caused by various stresses such as conventional chemotherapeutic agents, allows these cancer cells to escape DNA damage-induced cell death [22, 23].

ROS/RNS overproduction, deregulation of host DDR machinery, and chronic inflammation, are the most common features of viral persistent infections, and together with non-selective autophagy, have also been documented in latency of herpesviruses including Epstein-Barr Virus (EBV) [24–32]. Moreover, we and others have provided overwhelming evidence supporting that EBV latent infection reprograms the host ubiquitin machinery for its own benefits [33–35], including the employment of linear ubiquitin chain assembly complex (LUBAC)-mediated ubiquitination to modulate LMP1 signal transduction [36]. However, as the major selective autophagy receptor and a ubiquitin sensor, p62 and its relationship with EBV latency and oncogenesis have never been investigated. In our recent publication, our findings have implied a role for the p62-autophagy interplay in ROS-elicited DDR in EBV latency [37].

In this study, we aimed to investigate the potential role of p62-mediated selective autophagy in regulating DDR in EBV latent infection. Our results show that p62-mediated selective autophagy is constitutively induced in virus-transformed cells, and correlates with ROS-Keap1-NRF2 pathway activity, and that a well-balanced basal level of p62-mediated selective autophagy is essential for maintaining genomic stability in this setting.

## Results

### p62-mediated selective autophagy is constitutively induced in viral latency and correlates with ROS-Keap1-NRF2 pathway activity

Our recent findings have shown that treatment of EBV+ cells with the calcium ionophore ionomycin, which raises the intracellular level of calcium (Ca^2^+) essential for ROS production, elevates the protein levels of both p62 and LC3b-II (the smaller cleavage product of LC3b, which generally represents a marker of autophagosomal activity in mammalian cells), and induces DNA damage; autophagy deficiency also elevates p62 protein levels [37]. Since both p62 and LC3b are targeted by autophagy for degradation, their turnover represents the autophagic flux (autophagic degradation activity) [38, 39]. These results have implied that the p62-autophagy interplay may be involved in oxidative stress in EBV latent infection.

We thus sought to evaluate the correlation between intracellular ROS, and p62 and autophagy levels in EBV latency programs. Results show that, although nearly 100% cells of each tested virus-associated cancer cell line produce ROS, their levels, as indicated by mean of fluorescence intensity (MFI), are consistently higher in SavIII, JiJoye, and MT4, compared to SavI, P3HR1, and CEM, respectively (Fig 1, A). Correspondingly, the cell lines with higher ROS production remarkably express higher p62 consistently at both protein and mRNA levels (Fig 1, B and C). Furthermore, the basal levels of p62-mediated selective autophagy activity, as indicated by both the cleaved LC3b product LC3b-II and phosphorylation of p62(Ser403), correspond to the endogenous ROS levels, and are readily detectable by immunoblotting in EBV type III latency and human T-cell leukemia virus-1 (HTLV1)-transformed MT4 cell line (Fig 1, B). Phosphorylation of p62(Ser403), which promotes p62-Ub binding, is crucial for activation of p62-selective autophagy [40]. Furthermore, interaction between selective autophagy receptors and Ub-like proteins (UBLs), such as LC3b, is the molecular basis for selective autophagy [41, 42]. Our IP results show that endogenous p62 and LC3b interact in virus-transformed cells (Fig 1, D). These results indicate that a basal level of p62-mediated selective autophagy is constitutively induced and correlates with the endogenous ROS level in viral latency.

**Fig 1.**
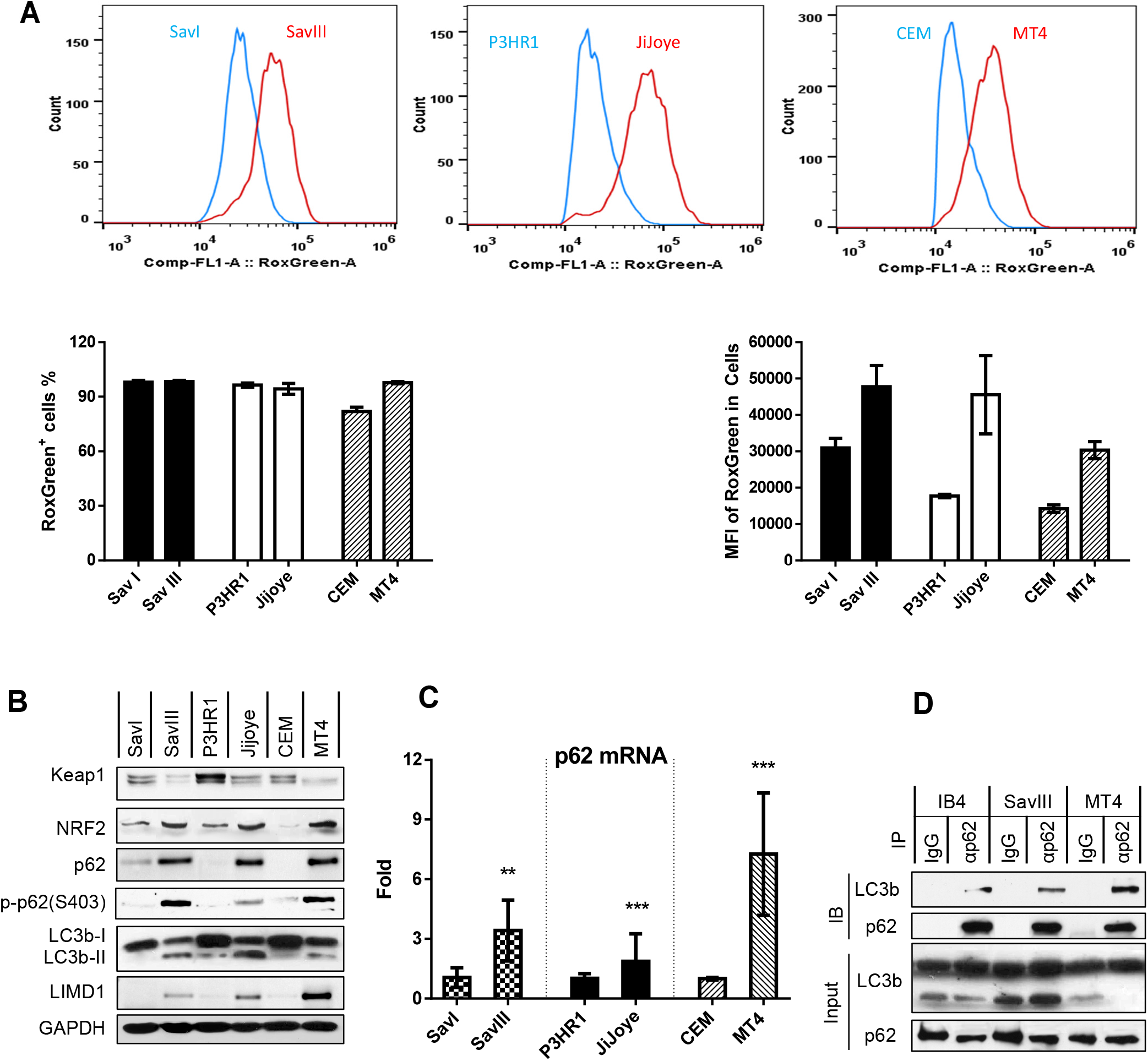
p62-mediated selective autophagy is endogenously induced in viral latency and correlates with ROS-Keap1-NRF2 pathway activity. **A**. Endogenous ROS production in viral latency was measured by flow cytometry with the CellRox Green reagent (Invitrogen). Three independent repeats were conducted, and representative results are shown. Results are the mean ± standard error (SE) of duplicates for each sample. MFI=mean fluorescence intensity. **B**. The correlation of endogenous p62 protein levels and autophagy activity with the Keap1-NRF2 pathway activity in viral latency was evaluated by immunoblotting with indicated antibodies. **C**. The correlation of p62 expression with viral latency was evaluated at the transcription level by qPCR. The mRNA levels in SavI, P3HR1, and CEM were set to 1, and compared with the paired SavIII, JiJoye, and MT4 cell lines, respectively. **D**. Interaction of endogenous p62 with LC3b in virus-transformed cells was evaluated by IP. Cell lysates (1 mg each) were pre-cleared with mouse IgG (Sigma) before subjected to IP with IgG or anti-p62 clone D-3 (Santa Cruz). immunoprecipitants and inputs (5% of cell lysates) were probed with indicated antibodies.

The antioxidant transcription factor NRF2 is spontaneously degraded by the ubiquitin (Ub) E3 ligase complex Keap1/Cul3/RBX1 under normoxia; ROS/oxidative stress triggers autophagic degradation of Keap1, resulting in the accumulation and activation of NRF2, which then induces p62 expression [43, 44]. Thus, we evaluated the Keap1-NRF2 pathway activities, indicated by Keap1 and NRF2 expression levels, in these cell lines. As indicated in Fig 1, B, the endogenous p62 levels positively correlate with the Keap-NRF2 pathway activities, strongly suggesting that the Keap1-NRF2 pathway induces p62 expression at least in part in viral latency. In support of our findings, Keap1-NRF2 pathway is also activated in KSHV latency [45, 46].

Together, these results indicate that p62-mediated selective autophagy is constitutively induced in oncovirus latency, and correlates with the endogenous ROS-Keap1-NRF2 pathway activity.

### ROS that correlate with Keap1-NRF2 pathway activity contribute to p62 expression and activation of p62-mediated autophagy in viral latency

We next aimed to verify the induction of p62 by the ROS-Keap1-NRF2 pathway in viral latency. To this end, we first treated SavI and SavIII cells with the clinical topoisomerase II inhibitor doxorubicin (Doxo), which generates the highest level of mitochondrial ROS causing DSBs [47]. Results show that Doxo augments the activities of the Keap1-NRF2 pathway in both cell lines in a time-dependent manner. Interestingly, p62 protein levels and p-p62(S403) are increased by Doxo treatment in SavI cells, but p62 protein levels are decreased in SavIII cells. Moreover, LC3b-II is increased in both cell lines but decreased at late stage in SavIII cells (Fig 2, A). These results are consistent with the notion that considerable levels of ROS are required for induction of p62 and mediated autophagy, but excess ROS and selective autophagy result in autophagic degradation of p62 and LC3b in that both are targets of p62-mediated selective autophagy [38, 39].

**Fig 2.**
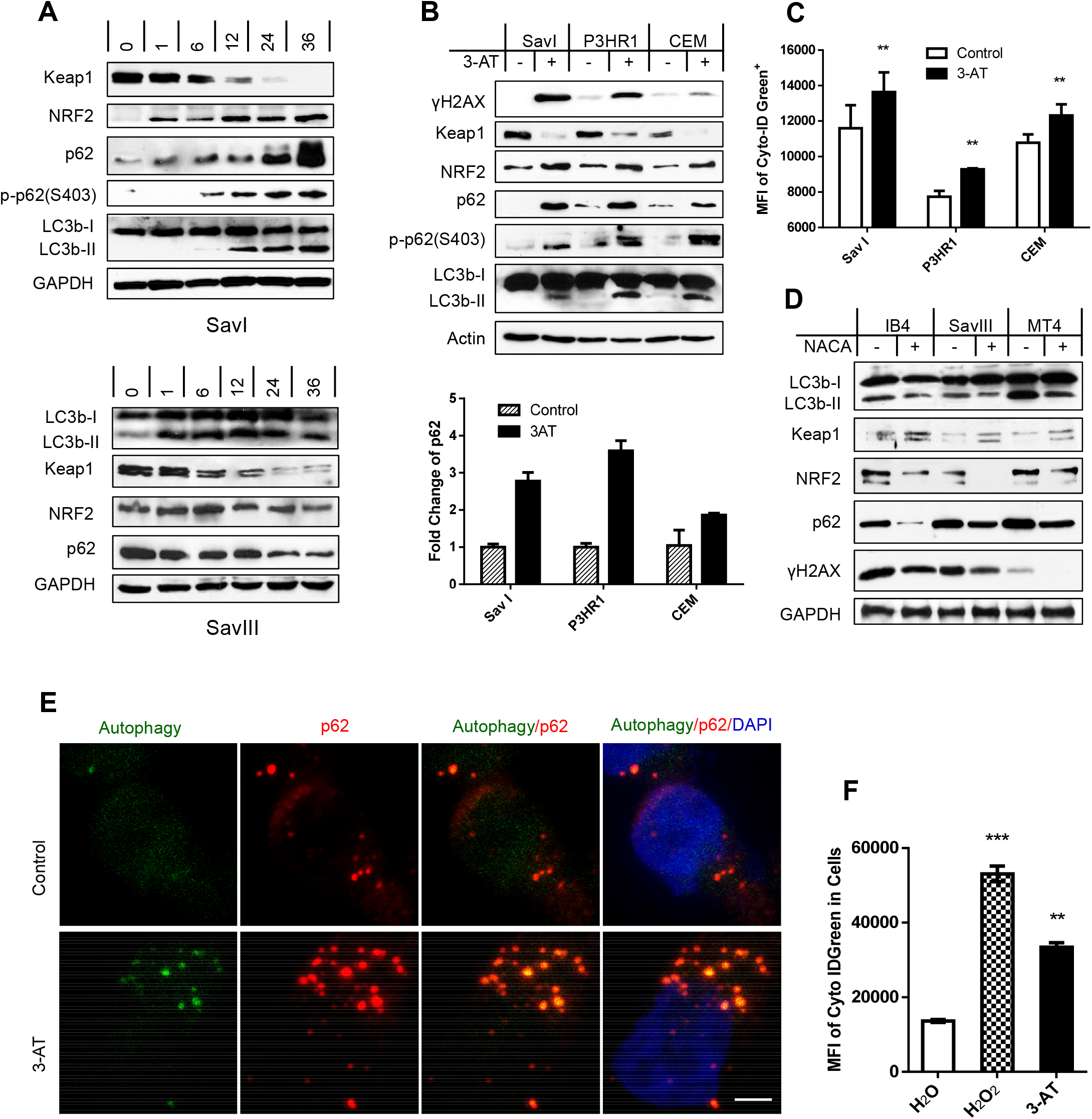
ROS correlate with Keap1-NRF2 pathway activity and contribute to p62 expression and activation of p62-mediated autophagy in viral latency. **A**. SavI and SavIII cells, which were derived from the same patient, were treated with 2 μM of the topoisomerase II inhibitor doxorubicin HCl (Doxo) (UBPBio) for different time periods. **B-C**. SavI, P3HR1, and CEM were treated with 20 mM of the H_2_O_2_ catalase 3-amino-1,2,4-triazole (3-AT) (Fisher Scientific) or vehicle control for 48 h. Cells were then subjected to IB, qPCR, and flow cytometry analyses. **D**. IB4, LCL45, and MT4 were treated with 3 mM of the antioxidant N-acetylcysteine amide (NACA) (Sigma) or vehicle control for 30 h. The treated cells were then subjected to analyses for p62, autophagy, Keap1-NRF2 pathway, and ROS production. **E**. IB4 cells were treated with 20 mM 3-AT (Fisher Scientific) or vehicle control for 48 h, and analyzed for p62-autophagosome colocalization by confocal microscopy. Living cells were first stained with a Cyto-ID Autophagy Detection kit (Enzo), and then fixed for staining with the mouse p62 antibody (D-3) and Alexa 555 coupled anti-mouse antibody (Invitrogen). Bar=2 μm. **F**. IB4 cells were treated with 50 μM H_2_O_2_ for 30 min and then medium was replaced and continued in culture for 2 days, or treated with 20 mM 3-AT for 48 h, before subjected to analysis of autophagy flux by flow cytometry with the Cyto-ID Autophagy Detection kit. MFI=mean fluorescence intensity.

We next used 3-amino-1,2,4-triazole (3-AT) to inhibit the endogenous activities of catalase, an enzyme converting H_2_O_2_ to H_2_O+O_2_, in cell lines with lower ROS levels to elevate their endogenous ROS levels. Then, we evaluated the Keap1-NRF2 pathway activity, and p62, autophagy, and DNA damage levels. Results show that 3-AT treatment substantially elevates endogenous levels of the Keap1-NRF2 pathway activities and p62 expression at both protein and mRNA levels, and also induces p62(S403) phosphorylation, autophagy and the DNA damage hallmark γH2AX that are readily detectable (Fig 2, B). Accumulation of LC3-II does not necessarily reflect an increased autophagic activity; instead it may represent its decreased clearance due to the blockage of autophagic degradation. Thus, we further measured autophagy flux, as indicated by MFI, by flow cytometry (Fig 2, C). In contrast, quenching endogenous ROS with the ROS scavenger N-acetylcysteine amide (NACA) in indicated cell lines substantially dampens p62 levels and autophagy activities due to blockage of their endogenous Keap1-NRF2 pathway activities, as well as attenuates endogenous DNA damage (Fig 2, D). Furthermore, confocal microscopy (Fig 2, E) and flow cytometry (Fig 2, F) results show that treatment of IB4 cells with 3-AT or with the traditional oxidative DNA damage inducer H_2_O_2_ remarkably increases p62 expression, autophagosomes and autophagy flux, and that most p62 foci co-localize with autophagosomal bodies in the cytoplasm.

Taken together, these results indicate that endogenous ROS, which correlate with Keap1-NRF2 pathway activity, are responsible for p62 expression and for induction of p62-mediated selective autophagy in viral latency.

### Autophagy inhibition sensitizes EBV^+^ cells to ROS-induced DNA damage that is associated with p62 accumulation

Since we have previously shown that mild ionomycin treatments induce p62 expression and profound autophagy in lymphoblastic cell lines (LCLs), but stringent treatments promote p62 degradation due to induction of massive autophagy [37], we used ionomycin treatment here to study the p62-autohagy interplay in regulating DDR in LCLs, which serve as a system crucial for genetic and functional study of carcinogen sensitivity and DNA repair [48].

We first used the lysosome-specific inhibitor bafilomycin A1 (BafA1), which inhibits lysosomal activity that occurs after LC3 processing, to inhibit autophagy activity induced by ionomycin in LCLs. Results show that both p62 and LC3b, which are both selectively degraded by p62-mediated autophagy [38, 39], are accumulated in ionomycin-treated cells due to impaired autophagy activities. As a consequence, the levels of γH2AX are remarkably augmented (Fig 3, A and B). To minimize the interference of potential “off-target” effects of BafA1, we performed this experiment using another lysosome-specific inhibitor chloroquine, and obtained similar results (Fig 3, C). These results indicate that the autophagy-p62 interplay plays a role in DDR in EBV latency.

**Fig 3.**
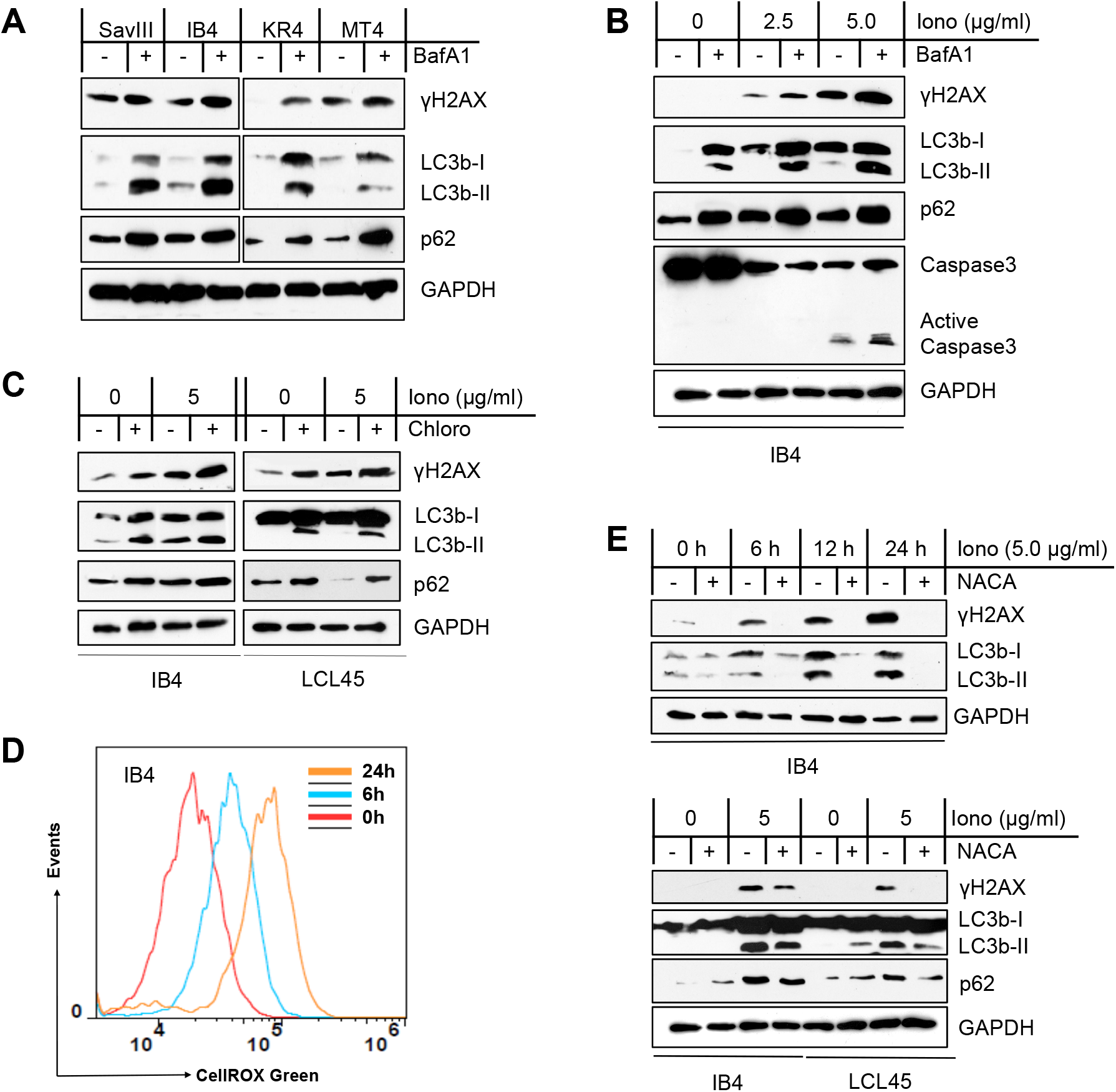
Autophagy inhibition sensitizes EBV+ cells to ROS-induced DNA damage that is associated with p62 accumulation. **A**. Cell lines with higher endogenous autophagy levels were treated with 0.4 μM of the vacuolar ATPase inhibitor bafilomycin A1 (BafA1) (Sigma) or vehicle control for 24 h, and then DNA damage (γH2AX) was evaluated by immunoblotting. **B-C**. The LCL lines IB4 and LCL45 were treated with ionomycin (Iono) (Sigma) with indicated concentrations for 48 h plus 0.4 μM BafA1 or vehicle control for 24 h or plus 50 μM of the lysosome inhibitor chloroquine (Chloro) (MP Biomedicals) or vehicle control for 6 h. p62, autophagy, and γH2AX were analyzed by immunoblotting. **D**. IB4 cells were treated with 5 μg/ml Iono or vehicle control for different time periods, and ROS production was measured by flow cytometry with the CellRox Green reagent (Invitrogen). A representative result from three independent repeats is shown. **E**. IB4 and LCL45 cells were treated with 5 μg/ml Iono plus 3 mM NACA or vehicle control for different time periods (upper panel) or for 30 h (lower panel). p62, autophagy, and γH2AX were analyzed by immunoblotting.

We further show that ionomycin triggers profound ROS production and DNA damage in LCLs in a time-dependent manner (Fig 3, D and E), and the ROS scavenger NACA offsets the effects of ionomycin (Fig 3, E). Thus, these results indicate that ROS are responsible for ionomycin-induced autophagy and DNA damage in EBV latency.

### Autophagy inhibition promotes DNA damage-induced cell death in association with p62 accumulation in the nucleus

To confirm the requirement of ROS for induction of autophagy and DNA damage in EBV latency, we further used H_2_O_2_ to treat IB4 cells. Results show that H_2_O_2_ treatment induces profound DNA damage and reduces the endogenous p62 protein level in a dose-dependent manner, and blockage of autophagy activity with BafA1 potentiates the DNA damage that correlates with elevated p62 protein levels (Fig 4, A).

**Fig 4.**
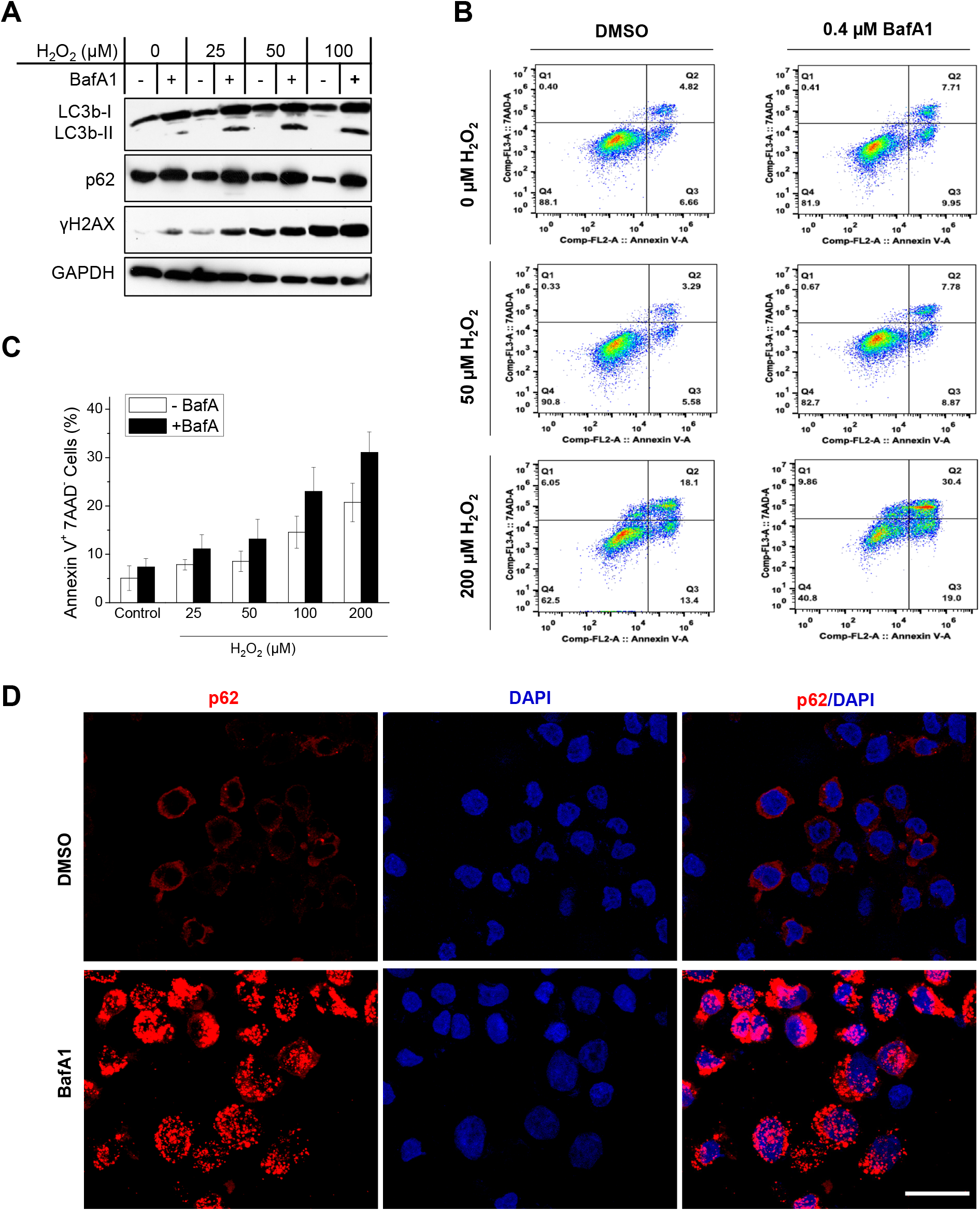
Autophagy inhibition promotes cell death in association with p62 accumulation in the nucleus. IB4 cells were treated with indicated concentrations of H_2_O_2_ in medium for 30 min. Then the medium was removed, and 0.4 μM BafA1 (Sigma) or vehicle control was added in freshly replaced medium for 48 h. **A**. p62, autophagy, and DNA damage were analyzed by immunoblotting. **B-C**. Cell death was analyzed by flow cytometry for Annexin V and 7-AAD expression. Results from a representative experiment of five independent repeats are shown (B), and statistical analysis results are expressed as mean ± standard error (SE) (C). **D**. p62 subcellular localization was visualized under confocal microscope. Bar=10 μm.

Furthermore, autophagy inhibition by BafA1 promotes cell death (as indicated by 7-AAD expression and Annexin-V binding, or caspase-3 activity) induced by H_2_O_2_ (Fig 4, B and C) or ionomycin (Fig 3, B), respectively. Importantly, confocal microscopy results further show that p62 translocates from the cytoplasm to the nucleus in response to autophagy inhibition (Fig 4, D).

Taken together, these results (Figs 3 and 4) indicate that autophagy inhibition exacerbates ROS-induced DNA damage by promoting p62 stabilization and nuclear translocation, further supporting that p62-mediated autophagy promotes DDR in EBV latency.

### Nuclear p62 accumulation upon autophagy inhibition destabilizes HR DNA repair proteins CHK1 and RAD51 in viral latency

It has been reported that autophagy inhibition or nuclear p62 accumulation accruing from autophagy deficiency promotes proteasomal degradation of HR DNA repair proteins such as RAD51, CHK1 and FLNA [17, 49]. Our results show that SavIII, JiJoye, MT4, and IB4, which have higher endogenous p62 and autophagy levels (Fig 1, B), have lower CHK1 and RAD51 protein levels as well as CHK1 activity (as indicated by phosphorylation of CHK1(S345)), compared to SavI, P3HR1, CEM, and BJAB, respectively (Fig 5, A). These results suggest that the p62-autophagy interplay may also regulate proteasome-dependent stability of CHK1 and RAD51 proteins and CHK1 activity in viral latency. It has also been reported that p62 promotes NHEJ by activating the Keap1-NRF2 pathway in a feedback loop, which consequently induces expression of NHEJ-specific repair proteins such as 53BP1 [50]. However, our results show that 53BP1 reversely correlates with p62 at the protein level in viral latency (Fig 5, A), indicating that NHEJ-mediated DNA repair activity is also compromised in virus-transformed cells. In addition, endogenous DNA damage (as indicated by γH2AX expression) is consistently lower in viral latency with higher p62-mediated autophagy levels, in which both HR and NHEJ pathways are deficient (Fig 5, A), supporting our hypothesis that p62-mediated autophagy functions as an alternative mechanism that enables these cells to resist to DNA damage.

**Fig 5.**
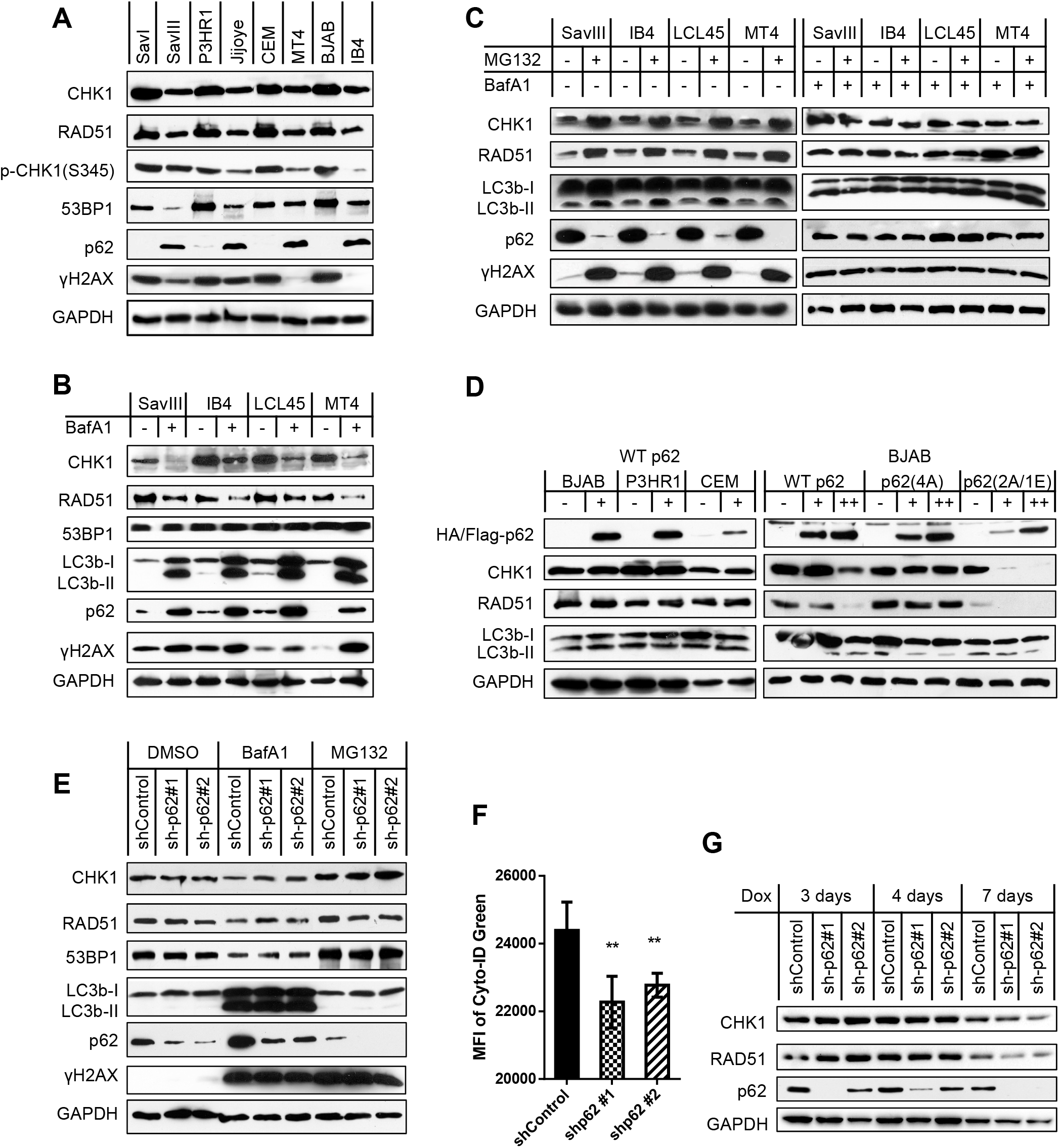
p62 accumulation upon autophagy inhibition destabilizes HR DNA repair proteins CHK1 and RAD51 in viral latency. **A**. Correlation of p62 with CHK1, RAD51 and 53BP1 protein levels in viral latency were analyzed in paired cell lines. **B**. Cell lines with higher levels of p62 protein were treated with 0.4 μM BafA1 (Sigma) for 48 h. Indicated proteins were probed by immunoblotting. **C**. Cell lines with higher levels of p62 protein were treated with 10 μM of the proteasome inhibitor MG132 for 6 h (left panel), or pre-treated with 0.4 μM BafA1 before MG132. Indicated proteins were probed by immunoblotting. **D**. Cell lines with lower levels of p62 protein were transfected with 5 μg (+) or 10 μg (++) of HA-p62 plasmids, its mutants with Flag tag, or vector control in each electroporation (1X10^7^ cells). Indicated proteins were analyzed by immunoblotting 48 h post-transfection. **E-F**. IB4 cells stably harboring p62 shRNA or shRNA control in 1 μg/ml puromycin were treated with 0.4 μM BafA1 for 48 h or 10 μM MG132 for 6 h, before subjected to immunoblotting or flow cytometry. shRNA expression was induced by 1 μg/ml doxycycline for 2 days before the drug treatments. MFI=mean fluorescence intensity. **G**. CHK1 and RAD51 protein stability was evaluated by immunoblotting in virus-transformed IB4 cells stably expressing control shRNA or p62 shRNA that were induced by 1 μg/ml doxycycline for different time points.

To check if autophagy has a role in regulation of the stability of these DNA repair proteins in virus-transformed cells, we inhibited endogenous autophagy activities with BafA1. Results show that autophagy inhibition did not affect 53BP1, but clearly decreases CHK1 and RAD51 protein levels that are associated with elevated endogenous p62 protein levels (Fig 5, B).

To validate whether proteasome also mediates degradation of CHK1 and RAD51 in virus-transformed cells, we used the proteasome inhibitor MG132 to treat these cells that express high levels of endogenous p62. As expected, our results show that MG132 treatment remarkably increases the protein levels of CHK1 and RAD51 (Fig 5C, left panel), confirming that CHK1 and RAD51 protein stability is controlled in a proteasome-dependent manner in virus-transformed cells. In addition to its ability to inhibit proteasomal activity, MG132 is capable of inducing autophagy in various cancer cells [51]. Our results show that MG132 also increases the LC3b cleavage product LC3b-II in virus-transformed cells, and dramatically reduces p62 protein levels (Fig 5C, left panel).

To further validate the role of autophagy-p62 interplay in proteasomal degradation of CHK1 and RAD51, we pre-treated the cells with BafA1 to inhibit autophagy before inhibition of proteasome activity. Results show that MG132 failed to cause CHK1 and RAD51 accumulation after p62 restoration by autophagy inhibition (Fig 5C, right panel). These observations are in line with the previous report showing that CHK1 and RAD51 accumulation is attributable to p62 depletion resulting from autophagy inhibition [18].

Together, our results indicate that the CHK1 and RAD51 protein levels reversely correlate with the p62 levels in viral latency, implying a role of the autophagy-p62 interplay in negative regulation of their proteasome-mediated stability. However, the accumulation of CHK1 and RAD51 and decrease of p62 after MG132 treatment did not mitigate DNA damage; instead, DNA damage is strikingly increased (Fig 5, C). This observation can be explained by the fact that proteasome function is required for DNA damage repair [52].

Next, we sought to evaluate whether transient expression of p62 regulates DNA repair protein stability. To this end, we transfected the expression plasmids harboring HA-p62 or pcDNA4 vector control into BJAB, P3HR1, and CEM cells, which express low levels of endogenous p62. Then, we analyzed CHK1, RAD51, and 53BP1. Surprisingly, results show that exogenic expression of p62 did not affect the protein levels of CHK1 and RAD51, and did not induce LC3b cleavage either (Fig 5D, left panel). By checking total p62 expression, we found that transfection of p62 expression plasmids did not evidently increase p62 levels, likely due to relatively low transfection efficiency of B cells (data not shown). To address this issue, we transfected these cells with more p62 plasmids, and results show that a greater p62 expression is able to downregulate these proteins (Fig 5D, right panel). We further employed two p62 mutants, with p62(4A) constitutively localizing in the cytoplasm and p62(2A/1E) mainly localizing in the nucleus, since a recent report shows that nuclear accumulation of p62 responding to autophagy inhibition is required for its degradation of CHK1 and RAD51 [18]. As expected, the nucleus-localizing mutant p62(2A/1E), but not the cytoplasm-localizing mutant p62(4A), remarkably decreases CHK1 and RAD51 protein levels (Fig 5D, right panel). Together with Fig 5, B and C, these results suggest that p62-mediated destabilization of DNA repair proteins is dependent on its accruing from autophagy inhibition, which we show results in p62 nuclear translocation (Fig 4, D).

To further confirm the conditional role of p62 in downregulation of the DNA repair proteins, we depleted p62 expression in IB4 cells using shRNA-mediated gene knockdown. Results show that p62 depletion reduces autophagy activity, as shown by decreased autophagy flux (Fig 5, F), consistent with the notion that p62 is required for endogenous autophagy induction. However, the protein levels of CHK1, RAD51, and 53BP1 have no consistent and apparent changes in p62-depleted cells (Fig 5, E and G). Additional inhibition of the residual autophagy activities, or treatment with MG132, which induces autophagy (Fig 5, C), also did not cause apparent difference on their levels in p62-depleted cells versus control cells (Fig 5, E). Considering that the majority of p62 is spontaneously located in the cytoplasm of virus-transformed cells (Fig 4, D), shRNA-mediated depletion might have minor effect on nuclear p62. Thus, these results are indeed consistent with our findings and a recent report that nuclear localization of p62 is required for destabilization of DNA repair proteins [18].

Collectively, these results demonstrate that p62 accumulation in the nucleus in response to autophagy inhibition promotes proteasomal degradation of RAD51 and CHK1 in virus-transformed cells. These results also indicate that a fine balance of p62 and autophagy levels is required to confine endogenous p62 in the cytoplasm of virus-transformed cells. Further study is required to determine how autophagy inhibition causes p62 accumulation in the nucleus.

### p62 depletion by RNA interference promotes RNF168-mediated chromatin ubiquitination and DNA repair in viral latency

It has been shown that p62 inhibits both HR and NHEJ [18], through its physical interaction with RNF168, which mediates histone ubiquitination that is prelude to activation of both HR and NHEJ DSB repair mechanisms [19].

Consistently, our confocal microscopy results show that RNF168 co-localizes with endogenous γH2AX DNA damage foci in IB4 cells. Our immunoblotting results also demonstrated that RNF168 did not change in protein level in BafA1-treated cells (not shown). More importantly, shRNA-mediated p62 depletion significantly increases RNF168-γH2AX foci (Fig 6, A and B). Further, using the anti-ubiquitinated proteins antibody FK2, we show that shRNA-mediated p62 depletion significantly increases ubiquitination in the nucleus and FK2-γH2AX foci (Fig 6, C and D). We further validated the interaction of endogenous p62 with RNF168 by immunoprecipitation (IP) in virus-transformed cells treated with BafA1 (Fig 6, E), and the increased histone H3 ubiquitination due to p62 depletion in cells treated with ionomycin (Fig 6, F). Our IP assays failed to detect definite p62-RNF168 interaction and H3 ubiquitination in these cells without treatments (not shown). In conclusion, these results indicate that p62 depletion promotes chromatin ubiquitination and RNF168-mediated DNA repair mechanisms.

**Fig 6.**
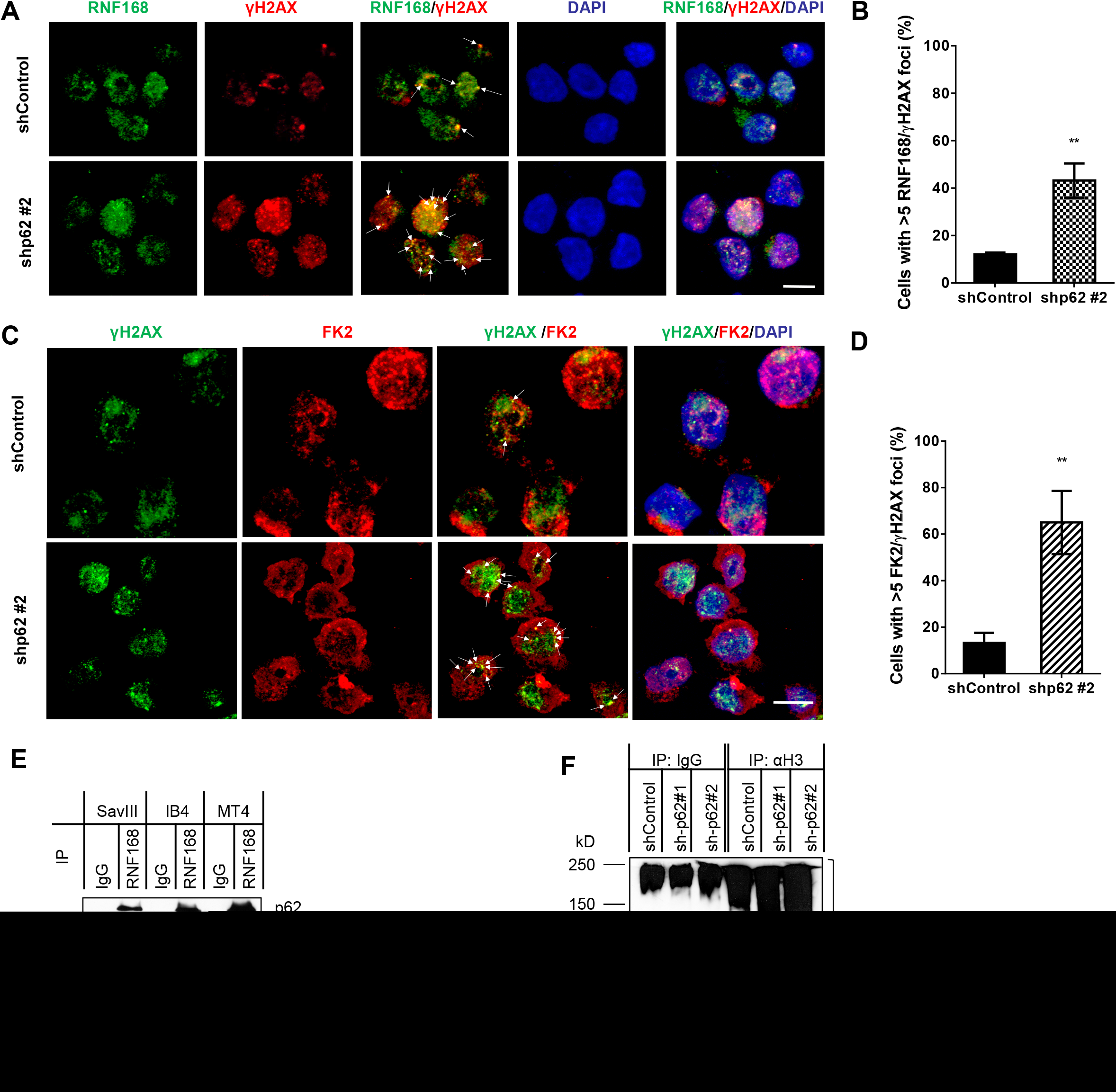
p62 depletion by RNA interference promotes RNF168-mediated chromatin ubiquitination and DNA repair in viral latency. IB4 cells stably expressing control or p62 shRNA were maintained with 1 μg/ml puromycin, and shRNA expression was induced by 1 μg/ml doxycycline for 3 days before subjected to confocal microscopy analysis for: **A**. RNF168, γH2AX and their colocalization in the nucleus with a rabbit RNF168 antibody (Millipore) and an Alexa 488 coupled anti-rabbit antibody (Invitrogen), and a mouse γH2AX(S139) antibody (BioLegend) and an Alexa 555 coupled anti-mouse antibody (BioLegend); **C**. chromatin ubiquitination, γH2AX and their colocalization in the nucleus with the mouse ubiquitin antibody clone FK2 (Millipore) and the Alexa 555 coupled anti-mouse antibody, and a rabbit γH2AX(S139) antibody (Cell Signaling Technol.) and the Alexa 488 coupled anti-rabbit antibody. Bar=6 μm. **B** and **D**. Quantification of RNF168/γH2AX foci or FK2/γH2AX foci in A and C, respectively. **E**. Interaction between RNF168 and p62 was assessed by immunoprecipitation. The indicated cell lines were treated with 0.4 μM BafA1 for 48h, and then cell lysates (0.5 mg total proteins for each) were subjected to immunoprecipitation with a rabbit RNF168 antibody (Proteintech Group Inc.), and immunoprecipitated proteins were analyzed by immunoblotting with the p62 antibody and the sheep RNF168 antibody (Invitrogen). **F**. Histone H3 ubiquitination was assessed in p62-depleted IB4 cells treated with 2.5 μg /ml ionomycin for 48 h. Cell lysates (0.5 mg total proteins for each) from IB4 cells stably expressing control or p62 shRNA were subjected to denatured immunoprecipitation with a rabbit H3 antibody (Invitrogen), and immunoprecipitated H3 was probed with the FK2 antibody and the mouse H3 antibody (1G1) (Santa Cruz).

## Discussion

In this study, we provide several lines of evidence that support a crucial role for p62-mediated autophagy in regulation of DDR in oncogenic virus latent infection. First, p62 is upregulated by ROS that correlates with the activity of the Keap1-NRF2 pathway, and considerable levels of p62-mediated selective autophagy are constitutively induced in this setting. Second, inhibition of autophagy in virus-transformed cells exacerbates ROS-induced DNA damage, and destabilizes the DNA repair proteins RAD51 and CHK1 in a manner depending on p62 accumulation in the nucleus; in contrast, excess autophagy induction promotes accumulation of the DNA repair proteins CHK1 and RAD51 that is associated with p62 degradation. Third, shRNA-mediated p62 depletion promotes RNF168-mediated chromatin ubiquitination and DNA repair in EBV latency. These findings have defined a crucial role for p62-mediated autophagy in regulation of DDR in viral latency (Fig 7).

**Fig 7.**
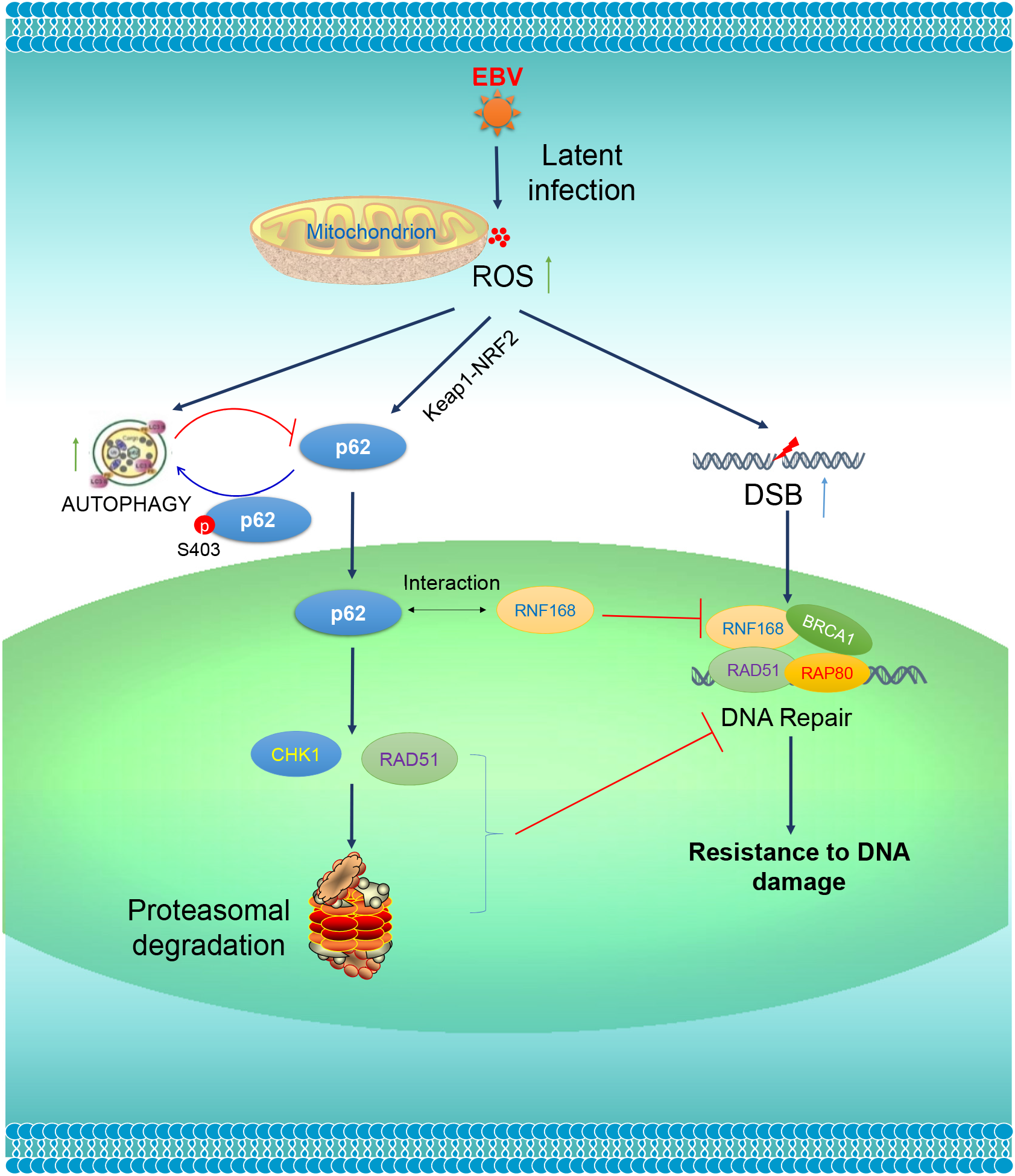
A diagram showing the interaction of p62-mediated autophagy with DDR in EBV latency. EBV latent infection produces ROS, which further induce p62 expression through the Keap1-NRF2 pathway and activate p62-mediated selective autophagy. ROS also cause DNA damage, including double strand breaks (DSBs). p62 accumulated in the nucleus due to autophagy inhibition inhibits DSB repair through promoting proteasome-mediated degradation of CHK1 and RAD51 and interacting with RNF168. A moderate level of endogenous p62-mediated autophagy in virus-transformed cells endows them with resistance to DNA damage. Loss of the p62-autophagy balance by exogenic stresses that inhibit autophagy or inducing ROS will exacerbate endogenous DNA damage.

The p62-autophagy interplay is well balanced and controlled in diverse contexts, with cancer and aging being two representative systems [4–6, 53]. Loss of this balance by exogenic or endogenous stresses may result in different impacts on DDR. Pharmaceutical inhibition of autophagy, or spontaneous autophagy deficiency during aging, chronic inflammation, or neurodegeneration, leads to p62 accumulation, consequently attenuating DNA repair that accounts for the etiology of age-related disorders [54, 55]. In contrast, substantial enhancement of basal levels of autophagy in cancer cells by anticancer chemotherapeutic drugs or by radiation therapy promotes p62 degradation, and consequently confers these cells resistance to DNA damage-induced cell death [23, 56, 57].

Moreover, our results indicate that the basal levels of p62-mediated autophagy are distinctly regulated in different EBV latency programs. ROS are produced separately by the EBV products LMP1, EBNA1/2, and EBERs, amongst which LMP1 induces predominant ROS [58–62]. In consistent, our results show that EBV type III latency produces a greater level of ROS compared with type I latency (Fig 1, A). Thus, cells with type III latency express higher endogenous levels of p62-mediated autophagy (Fig 1, B), which are required to overcome the higher risk of DNA damage in response to endogenous higher oxidative stress and replication stress to support their aggressive proliferation, and growth and survival demands. As such, a higher level of p62-mediated autophagy also confers these cells greater resistance to DNA damage in response to drug treatments. The type III latency cell line P3HR1, which was derived from the parental JiJoye but lacks LMP1 expression, resembles type I latency cell lines in ROS production, and expression of p62 and autophagy (Fig 1), and DNA repair proteins (Fig 5), further supporting that LMP1 contributes the majority to these events.

Surprisingly, our results consistently show that, in contrast to p62 elimination by massive autophagy, p62 depletion by shRNA potentiates, but not alleviates, DNA damage (not shown). There are a few possibilities to explain this paradox. First, p62 depletion impairs p62-mediated selective autophagy that is resident in the cytoplasm (Fig 2, E) and required for maintaining the ability of the cell to resist to DNA damage (Fig 5, E and F); second, other p62-mediated, autophagy-independent, DNA damage-protecting functions are abrogated after shRNA-mediated p62 depletion and consequently DNA damage is accelerated, given the fact that p62 is a multifunctional protein [3]. Investigation of these potential p62-mediated but autophagy-independent functions in viral latency and oncogenesis is underway. In fact, p62 depletion resulting in worsen DNA damage is coincident with its role as a tumor promoter, which is induced by Ras that accounts for at least 25% of human cancers [63]. p62 overexpression in hepatocellular carcinoma (HCC) predicts poor prognosis [64].

Based on the observations from us and other relevant studies, we propose that p62 plays a dichotomous role in DDR, depending on the presence or absence of autophagy that determines the p62 protein level and subcellular localization. Higher levels of nuclear p62 resulting from defective autophagy inhibit DNA repair and therefore perturb genomic instability that facilitates tumor initiation [65]. In line with our findings, it has been reported that p62 ablation decreases tumorigenesis in mouse models with defective autophagy [63]. A recent report has also shown that autophagy resulting from telomere shortening during replicative crisis protects genomic stability, and acts as a suppressor of tumor initiation [66]. In contrast, considerable levels of p62 in cancer cells promote DNA repair by mediating selective autophagy activation in the cytoplasm, and consequently confer these cancer cells resistance to DNA damage. p62 is upregulated at considerable levels in different cancer cells, including breast and prostate cancers, where it is required for induction of selective autophagy to support cancer cell metabolism and survival [64, 67, 68].

Regarding the mechanisms underneath deregulation of DDR by the p62-autophagy interplay, our results indicate that autophagy inhibition promotes proteasomal degradation of RAD51 and CHK1 in a manner depending on p62 accumulation in the nucleus (Fig 5), and that p62 depletion promotes RNF168-mediated DDR (Fig 6). Our results show that the majority of endogenous p62 is tethered to autophagosomes in the cytoplasm (Fig 2, E), but inhibition of DNA repair requires p62 in the nucleus [18]. Consistently, we show that autophagy inhibition promotes p62 nuclear translocation (Fig 4, D), although the mechanism remains to be disclosed. Thus, our results define a new role for p62-mediated autophagy in preventing DNA damage by confining p62 in the cytoplasm. In conclusion, p62-mediated selective autophagy not only confers invulnerability to DNA damage, but also at least partially contributes to the deficiency of traditional DDR mechanisms, in virus-transformed cells. In this regard, it is to our understanding that an oncogenic virus gains a dual benefit by invoking p62-mediated autophagy: one facet of p62-mediated autophagy endows its host cell with ability to resist DNA damage to support cell survival; the other facet hijacks the traditional DDR mechanisms in the host cell to facilitate genomic instability.

Although p62 was reported to promote NHEJ by inducing 53BP1 expression through the Keap1-NRF2 pathway that requests p62 S349 phosphorylation [50, 69], our results show that p62 reversely correlates with 53BP1 in viral latency (Fig 5, A), and that p62 accumulation by autophagy inhibition (Fig 5, B), transient expression of p62, or p62 depletion by shRNA (not shown), failed to regulate 53BP1 levels. Thus, p62 does not regulate 53BP1 in our system, and how 53BP1 is downregulated in virus-transformed cells is worthy of further examination. Rather, p62 interacts with RNF168 in response to autophagy inhibition, and consequently inhibits RNF168-mediated chromatin ubiquitination in viral latency (Fig 6). In this regard, our findings are consistent with a recent study [18], which has shown that p62 can impede both HR and NHEJ through its interaction with RNF168, given that RNF168-mediated histone ubiquitination is prerequisite for activation of all DSB repair mechanisms [19].

Viruses have evolved diverse strategies to hijack host traditional DDR machinery during their chronic infections to perturb genomic integrity, including their ability to deregulate the p62-autophagy balance, which we believe only makes partial contribution. In fact, our group has recently shown that traditional DDR mechanisms are also deficient in aging T cells in chronic HCV infection [70, 71], at least partially attributable to endogenous p62 accumulation accruing from deficient autophagy in these cells. p62 also inhibits DDR through other mechanisms that have not been fully elucidated. For example, nuclear p62 interacts with and inhibits PML nuclear bodies, which are involved in DNA repair [72, 73]. Moreover, other autophagy mechanisms, such as chaperone-mediated autophagy [74], also participate in DDR, by regulating stability of DDR-related proteins such as HP1α and CHK1 [75, 76], and by regulating p62-dependent or -independent cellular functions [77]. It is of great interest to investigate these potential mechanisms and their coupled cellular mechanisms in virus-mediated oncogenesis.

Endogenous ROS/RNS trigger signal cascades that activate both DDR and autophagy programs. Unrepaired damaged DNA can serve as a major source of genomic instability particularly in cancer cells where traditional DDR and cell death pathways are compromised.

Thus, cancer cells heavily rely on autophagy, not only to replenish their deficient DNA repair mechanisms, but also to corroborate their higher metabolic demand than normal cells do. Therefore, cancer cells are more vulnerable to autophagy inhibition, providing viable opportunities for therapeutic strategy by targeting autophagy, in particular in combination with another cellular mechanism that is specifically coupled with autophagy in a given cancer context for improving clinical efficacy and specificity [3, 20, 78].

## Materials and Methods

### Cell Lines

SavI, SavIII, P3HR1 and JiJoye are human B cell lines derived from EBV-positive Burkitt’s lymphoma (BL) patients. P3HR1 was derived from JiJoye but does not express LMP1 due to lacking the entire EBNA2 ORF in the viral genome [79]. BJAB is an EBV-negative BL line. The lymphoblastic cell line (LCL) IB4 was derived from umbilical cord B-lymphocytes latently infected with EBV *in vitro.* KR4 is a LCL with gamma irradiation resistance [80]. LCL45 is a newly established LCL by *in vitro* transforming primary B cells of a healthy adult peripheral blood with the EBV strain B95.8. CEM is a HTLV1-negative, EBV-negative T cell line derived from acute leukemia, and MT4 is a HTLV1-transformed CD4+ T cell line derived from umbilical cord blood lymphocytes. B and T cell lines are cultured with RPMI1640 medium plus 10% FBS and antibiotics. All cell culture supplies were purchased from Life Technologies.

### Antibodies and Reagents

p62 (D-3), LIMD1 (H-4), and histone H3 (1G1) mouse monoclonal antibodies were from Santa Cruz for immunoprecipitation or immunoblotting. p62-Alexa Fluor 488 mouse antibody for flow cytometry was from Millipore or R&D Systems. Phospho-p62(S403), NRF2 (D1Z9C) mouse monoclonal antibody, and HRP-coupled secondary antibodies were from Cell Signaling Technologies. RNF168 rabbit polyclonal antibody and the FK2 mouse monoclonal antibody that recognizes K29-, K48-, and K63-linked polyubiquitin chains and monoubiquitin conjugation but not free ubiquitin for immunofluorescence were from Millipore. The RNF168 rabbit polyclonal antibody for IP was from Proteintech Group Inc. RNF168 sheep polyclonal, and LC3b and histone H3 rabbit polyclonal antibodies were from Invitrogen. RAD51 rabbit polyclonal and CHK1 (G-4) mouse monoclonal antibodies were from Abcam and Santa Cruz, respectively. The γH2AX(S139) mouse and rabbit antibodies were from BioLegend and Cell Signaling Technologies, respectively. Mouse HA (clone HA-7) and Flag (clone M2) antibodies were from Sigma. Secondary antibodies coupled with FITC, Alexa Fluor, APC, PE, Cy5, or PerCP and human anti-CD19-PE were from BioLegend, BD Biosciences, Invitrogen, or eBioscience.

HA-p62 cloned in pcDNA4 was a gift from Yu-Ying He [81], and Flag-p62 mutants were gifts from Dr. Ying Zhao [18]. Flag-p62(R186A/K187A/K264A/R265A) (designated as p62(4A)) cannot localize in the nucleus and Flag-p62(K7A/D69A/I314E) (designated as p62(2A/1E)) mainly localizes in the nucleus [18]. CellRox Green, MG132, chloroquine, and doxorubicin HCl, were purchased from Invitrogen, EMD Millipore, MP Biomedicals, and UBPBio, respectively. Ionomycin calcium salt, N-acetylcysteine amide, bafilomycin A1, doxycycline, and mouse and rabbit IgG were purchased from Sigma. H_2_O_2_ was from Santa Cruz.

### Transfection and Selection of Stable Transfectants

A set of p62 shRNA cloned in pTRIPz/Puro comprising three individual p62 shRNA and a scramble control plasmids were purchased from Dharmacon. We selected two of them for this study and the targeting sequences on the human p62/SQSTM1 gene are: shRNA#1: 5’-TCTCTTTAATGTAGATTCG-3’ and shRNA#2: 5’-TCAGGAAATTCACACTCGG-3’. Lentiviral packing, preparation, infection, and selection of stable cells by puromycin (0.5 μg/ml) were performed as detailed in our previous publication [37]. shRNA expression was induced by 1 μg/ml doxycycline for 3 days, and then cells (expressing red fluorescence) were subjected to a second selection by FACS on a FACS/Aria Fusion Cell Sorter (BD Biosciences). Stable transfectants were maintained in complete medium plus 1.0 μg/ml puromycin.

For transfection of BJAB, P3HR1, and CEM, GenePulser XCell (Bio-Rad) was used with optimal programs. These representative cell lines were chosen in that they are easier to be transfected with this technique, compared with other B cell lines.

### Confocal Microscopy

Cells were fixed in 2% paraformaldehyde (PFA) for 20 min, permeabilized with 0.3% Triton X-100 in phosphate-buffered saline (PBS) for 10 min, blocked with 5% bovine serum albumin (BSA) in PBS for 1 h, and then incubated with indicated primary antibodies at 4 °C overnight. Cells were washed with PBS with 0.1% Tween-20 for three times, and then incubated with corresponding secondary antibodies coupled with FITC, Alexa Fluor, APC, PE, Cy5, or PerCP, at room temperature for 1 h. Cells were then washed and mounted with DAPI Fluoromount-G (SouthernBiotech, Birmingham, AL). Images were acquired with a confocal laser-scanning inverted microscope (Leica Confocal, Model TCS sp8, Germany).

### Immunoprecipitation and Immunoblotting

To assess endogenous p62-RNF168 interaction, 1×10^7^ cells for each sample were lysed with NP40 lysis buffer (150 mM NaCl, 1% NP-40, 50 mM Tris-pH 8.0, plus protease inhibitors), and cell lysates were subjected to immunoprecipitation (IP) with 1.5 μg anti-RNF168 for overnight, and then incubated with 40 μl Protein A/G beads (Santa Cruz) for 1 h. After three washes, proteins on beads were denatured in 1% SDS before subjected to immunoblotting (IB). IB was carried out with indicated antibodies and signals were detected with an enhanced chemiluminescence (ECL) kit following the manufacturer’s protocol (Amersham Pharmacia Biotech). For H3 ubiquitination assay, endogenous H3 was pulled down with an H3 antibody in denaturing IP, as detailed in our publication [82].

### RNA Extraction and Real-time Quantitative PCR

Total RNA was isolated from tested cells using an RNeasy Mini kit according to the manufacturer’s protocols (Qiagen). The eluted RNA was subjected to reverse transcriptase reactions, which were performed with the use of GoScript RT kit following the manufacturer’s instructions (Promega).

Quantitative real-time PCR (qPCR) was performed with the use of SYBR Green (Applied Biosystems), on a CFX96^TM^ Real-time PCR Detection System (Bio-Rad). All reactions were run in triplicates. Mean cycle threshold (Ct) values were normalized to 18s rRNA, yielding a normalized Ct (ΔCt). ΔΔCt value was calculated by subtracting respective control from the ΔCt, and expression level was then calculated by 2 raised to the power of respective -ΔΔCt value. The averages of 2^(-ΔΔCt) in the control samples were set to 1 or 100%. Results are the average ± standard error (SE) of triplicates for each sample. Primers for real-time qPCR are as follows: p62: F: 5’-CAGGCGCACTACCGCGATG-3’ and R: 5’-ACACAAGTCGTAGTCTGGGCAGAC-3’. Keap1: F: 5’-CCATGGGCGAGAAGTGTGTCC-3’; R: 5’-ACAGGTTGAAGAACTCCTCTTGCTTG-3’. 18s rRNA: F: 5’-GGCCCTGTAATTGGAATGAGTC-3’ and R: 5’-CCAAGATCCAACTACGAGCTT-3’.

### Flow Cytometry

Samples were fixed with 2% PFA for 20 min at RT, then wash with flow buffer (eBioscience). Samples were then incubated with PE-conjugated anti-human CD19 antibody (eBioscience) or isotype controls for 20 min at RT, then wash with flow buffer, followed by incubation with p62-Alexa Fluor 488 antibody for 60 min at RT. Samples were then washed with flow buffer, and analyzed with BD C6 plus flow cytometer.

For intracellular ROS measurement, 1X10^6^ cells in 500 μl medium per well were seeded in 24-well plates, and cultured overnight. 1 μl CellROX Green Reagent (Invitrogen) was added to each well and incubated for 30 min. Cells were then washed 3 times with PBS, and fixed with 2% PFA for 20min at RT, followed by extensive washes and then incubated with PE-conjugated anti-human CD19 antibody (eBioscience) for 20min at RT, before subjected to flow cytometry.

### Apoptosis Assays

Apoptosis was quantified using flow cytometry as detailed in our previous publication [37], for Annex V binding (BD Biosciences) and 7-Aminoactinomycin D (7-AAD) expression (eBioscience). Caspase 3 activity was evaluated by Western blotting.

### Statistical Analysis

Unpaired, two-tailed student *t* tests were executed using GraphPad Prism (version 6) to determine the differences between two data sets obtained from three independent experiments. p<0.05 (*) and p<0.01 (**), and p<0.001 (***) were considered significant. Data are expressed as mean ± standard error (SE) of duplicate or triplicate samples, and representative results from at least three independent repeats with similar results are shown.

## Acknowledgements

We thank Drs. Yu-Ying He and Ying Zhao for the p62 expression plasmids. This study was conducted with resources and the use of facilities at the James H. Quillen Veterans Affairs Medical Center. The contents in this publication are solely the responsibility of the authors and does not necessarily represent the official views of the NIH, the Department of Veterans Affairs, or the United States Government. The authors declare no competing interests.

